# Real-time library search increases cross-link identification depth across all levels of sample complexity

**DOI:** 10.1101/2022.11.16.516769

**Authors:** Max Ruwolt, Yi He, Diogo Borges Lima, William Barshop, Johannes Broichhagen, Romain Huguet, Rosa Viner, Fan Liu

## Abstract

Cross-linking mass spectrometry (XL-MS) is a universal tool for probing structural dynamics and protein-protein interactions *in vitro* and *in vivo*. Although cross-linked peptides are naturally less abundant than their unlinked counterparts, recent experimental advances improved cross-link identification by enriching the cross-linker modified peptides chemically with the use of enrichable cross-linkers. However, mono-links (i.e., peptides modified with a hydrolyzed cross-linker) still hinder efficient cross-link identification since a large proportion of measurement time is spent on their MS2 acquisition. Currently, cross-links and mono-links cannot be separated by sample preparation techniques or chromatography because they are chemically almost identical. Here, we found that based on the intensity ratios of four diagnostic peaks when using PhoX/tBu-PhoX cross-linkers, cross-links and mono-links can be partially distinguished. Harnessing their characteristic intensity ratios for real-time library search (RTLS)-based triggering of high-resolution MS2 scans increased the number of cross-link identifications from both single protein samples and intact *E. coli* cells. Specifically, RTLS improves cross-link identification from unenriched samples and short gradients, emphasizing its advantages in high-throughput approaches and when instrument time or sample amount is limited.

## Introduction

Cross-linking mass spectrometry (XL-MS) is a versatile tool that allows mapping of stable and transient protein-protein interactions (PPIs) in complex biological systems^1^. Moreover, it offers structural information on single proteins and protein complexes, thereby complementing structural biological techniques of higher resolution such as X-Ray crystallography and cryo-electron microscopy^2^. In XL-MS, the structural information is derived as distance restraints given by the length of the cross-linker and of the specific amino acid side chains it targets. The most commonly used cross-linkers react via NHS chemistry with primary amines (e.g., lysine residues) of one protein (intra-link), which gives insights into protein conformation, or two different proteins (inter-link), which sheds light on protein complex assemblies and PPIs. The depth of XL-MS coverage for both structural characterization and interactome profiling is highly dependent on the number of identified cross-links. Recent studies have put considerable efforts in the optimization of LC-MS parameters (e.g., charge filter, dynamic exclusion, fragmentation energy, and duty cycles)^3-5^, search engines^6-8^ and cross-linker designs^9-11^. For instance, cross-linkers decorated with an enrichment-handle allow removal of linear peptides, which are thought to make up 95 – 99% of the mixture. Enrichable cross-linkers greatly improve identification of low-abundant cross-links and lead to obviation of extensive fractionation^9^. Other approaches sought to take advantage of cross-linker chemistry by identifying fragmentation patterns unique to linear peptides, mono-links, and cross-links^12^. In the MS2 event of a mono- or cross-link, fragmentation of an amide bond occurs, either in the peptide backbone or the generated cross-link bond. Fragmentation at the peptide backbone leads to the formation of a tetrahydropyridine ring by carbon monoxide extrusion, followed by a loss of ammonia at the alpha carbon. Products of this reaction result in cross-linker specific diagnostic peaks for Lys mono-links and Lys-Lys cross-links^13^. These cross-linker specific fragments have proven useful to increase the speed and confidence of database searching^14^.

When using XL-MS for structural modeling, information from cross-links, mono-links and loop-links (formed when both ends of the cross-linker react with residues within the same peptide) is helpful to interpret solvent accessible surfaces and overall protein conformation. However, for large-scale interactomic studies, focusing on cross-links alone may be more beneficial as only cross-links provide information on residue-to-residue connections. Excluding unwanted species has recently been achieved in shotgun proteomics by applying real-time library search (RTLS). RTLS reduces acquisition time, increases quantification accuracy, and improves duty cycles^15, 16^ by raising measurement time for the ion species of interest. The ability of RTLS to focus MS data acquisition on precious, perhaps less abundant species makes it a promising approach for cross-link identification, but the full potential of RTLS-assisted XL-MS remains to be explored.

We recently described a sample processing workflow, in particular a two-step Immobilized Metal Affinity Chromatography (IMAC) enrichment strategy for interactome profiling of intact cells based on the enrichable cross-linker *tert*-butyl-PhoX (tBu-PhoX)^17^. In this study, we further augment the tBu-PhoX XL-MS pipeline from the data acquisition perspective by integrating RTLS to advance cross-link detection. We optimized instrument parameters for a fast MS2 survey scan that is searched against a custom-tailored library containing intensity ratios of diagnostic peaks from cross-links and mono-links. Based on the outcome of the search, potential cross-links trigger sequential high-resolution identification scans. We show that the RTLS-based acquisition strategy successfully boosts cross-link identifications, thereby providing a convenient tool for project guided acquisition in XL-MS measurements.

## Materials and Methods

### Cross-linking, protein digestion and IMAC enrichment

*E. coli* DH3α cells were grown in LB media for 16 h at 37 °C. Cells were pelleted at 800xg for 10 min at 4 °C. The cell pellet, chicken ovotransferrin, bovine serum albumin or yeast alcohol dehydrogenase (Sigma Aldrich) were resuspended in 20 mM HEPES pH 7.4 to a concentration of 10 mg/mL. 2.5 – 5 mM tBu-PhoX (Thermo Fisher Scientific) in DMSO was added to each solution and incubated for 30 – 60 min at room temperature. The cross-linking reaction was quenched by adding 20 mM Tris pH 8 and incubated for 15 min at room temperature. Cross-linked *E. coli* were incubated for 10 min at 95 °C after addition of 4% wt/v SDS and lyzed by sonication for 10 min using a Bioruptor (30 s cycles). *E. coli* proteins were precipitated by chloroform-methanol precipitation. Cross-linked proteins were supplemented with 8 M urea, reduced, alkylated and proteolyzed with Lys-C endopeptidase (1:200 wt/wt) and trypsin (1:100 wt/wt). The digestion was stopped after 16 h with 1% formic acid (FA). Cross-linked peptides were desalted using Sep-Pak C8 cartridges (Waters) and dried. Cross-linked and mono-linked peptides were enriched as described before^17^.

### LC-MS data acquisition

Samples were separated by reverse phase-HPLC using a Thermo Scientific™ EASY-nLC™ 1200 system connected to an EASY-Spray™ PepMap™ RSLC C18 column (0.075 mm x 250 mm, 2 μm particle size, 100 Å pore size (Thermo Fisher Scientific)) at 300 nL/min flow rate. The crosslinked samples were analyzed on the Orbitrap Eclipse™ Tribrid™ mass spectrometer with Instrument Control Software version 4.0. Reverse phase separation was accomplished using a 60 min or 125 min separation gradient (plus 10 – 20 min equilibration phase). Separation gradient: 3 – 40% solvent B (A: 0.1% FA; B: 80% ACN, 0.1% FA; Table S1). Cross-linked samples were analyzed using an HCD-MS2 acquisition strategy with 30% normalized collision energy (NCE) or stepped collision energy (SCE) 26 ± 5%. MS1 and MS2 scans were acquired in the orbitrap with a respective mass resolution of 60,000 and 30,000. Dynamic exclusion was set to 60 sec. Survey scans were acquired in the ion trap or the orbitrap with a scan range of m/z 120 – 500. For survey scans, precursor isolation window was set to 1.2 m/z and the normalized collision energy was determined experimentally. RTLS was performed by comparing survey scans with the spectral library entries representing the mono-link and cross-link classes in the “Reverse Library Search” mode with “Similarity Search” enabled. The library was implemented as described before^15^ and contained two synthetic spectra consisting of the four diagnostic peaks (m/z 201.1231, 215.1387, 312.0632, 377.1269) with their expected relative intensities. The annotation of the class within the spectral library was used to create instrument methods that either “reject” or “promote” the acquisition of additional scans for the desired class. Essentially, “promoting” a species forces its high-resolution MS2 acquisition regardless of the classification score (cosine score) while “rejecting” prohibits its acquisition. If none of the modes is chosen for the available classes, an individually adjusted cosine score threshold-based triggering will be performed. MS2 sequencing events were triggered for precursors at charge states +3 to +8.

### Data analysis

Spectral raw data files were analyzed using Proteome Discoverer 3.0 software (Thermo Fisher Scientific) with XlinkX node 3.0 using the noncleavable or noncleavable open search algorithms for cross-linked peptides and SEQUEST HT search engine for unmodified peptides and loop-links/mono-links. MS1 ion mass tolerance: 10 ppm; MS2 ion mass tolerance: 20 ppm. Maximal number of missed cleavages: 2; minimum peptide length: 6; max. modifications: 4; peptide mass: 500 – 8,000 Da. Carbamidomethylation (+57.021 Da) of cysteines was used as a static modification. PhoX cross-linked mass modifications were used as variable modifications for lysines or protein N-termini in addition to methionine oxidation (+15.995 Da). Data were searched for cross-links against a protein database containing chicken ovotransferrin (P02789), yeast alcohol dehydrogenase (P00330), bovine cytochrome C (P62894), bovine serum albumin (P02769), and proteases or a database generated from 714 protein identifications using the *E. coli* proteome retrieved from Uniprot as search space. The false discovery rate (FDR) was set to 1% at CSM and cross-link levels. Cross-linking analysis was also performed using pLink2^6^ (version 2.3.9). Search parameters were similar to those used in XlinkX except that the full *E. coli* proteome was used as search space. Post-processing and visualization were carried out using the rawrr R package^18^ and the XMAS plug-in^19^ for ChimeraX^20^.

## Results

The use of real-time library search for the exclusion of a specific pool of species such as mono-links requires a set of peaks that occur with high frequency in a stable intensity ratio to each other within the target species. To identify potential diagnostic peaks for mono-links and cross-links, we examined the peaks with the highest occurrence in 50,637 MS2 spectra from a previously published tBu-PhoX cross-linked *E. coli* dataset^17^ (Figure 1A). We observed four peaks (m/z 201.12, 215.12, 312.06, 377.13) in the low mass-to-charge region that appear in more than 50% of cross-linked and mono-linked spectra. The origin of peptide-independent cross-linker fragment ions m/z 312.06 and 377.13 (Figure 1C) was described before for cross-linkers of similar functionality, such as disuccinimidyl suberate (DSS) and disuccinimidyl sulfoxide (DSSO)^12-14^. Interestingly, certain peaks (e.g. m/z 201.12, 215.12, 243.13) that are observed for tBu-PhoX are also found with high occurrence in DSS data, suggesting that they are cross-linker-independent fragments (Figure S1). Thus, the four proposed diagnostic peaks (m/z 201.1231, 215.1387, 312.0632, 377.1269) are suited for the generation of a library, because they represent abundantly detected fragments specific for either cross-links or mono-links and cross-linker-independent species, ensuring an RTLS-based decision on almost all MS2 spectra. Assessing the relative intensity ratios of the chosen peaks showed differences of three m/z 312.06 containing ratios between cross-links and mono-links. While m/z 312 (z = 1) was ten times more intense in mono-links than in cross-links, the other three peaks showed similar and unaltered intensities between the two species (Figure 1B). We therefore considered these four peaks valuable for creating a library to exclude mono-links by RTLS.

**Figure 1.**
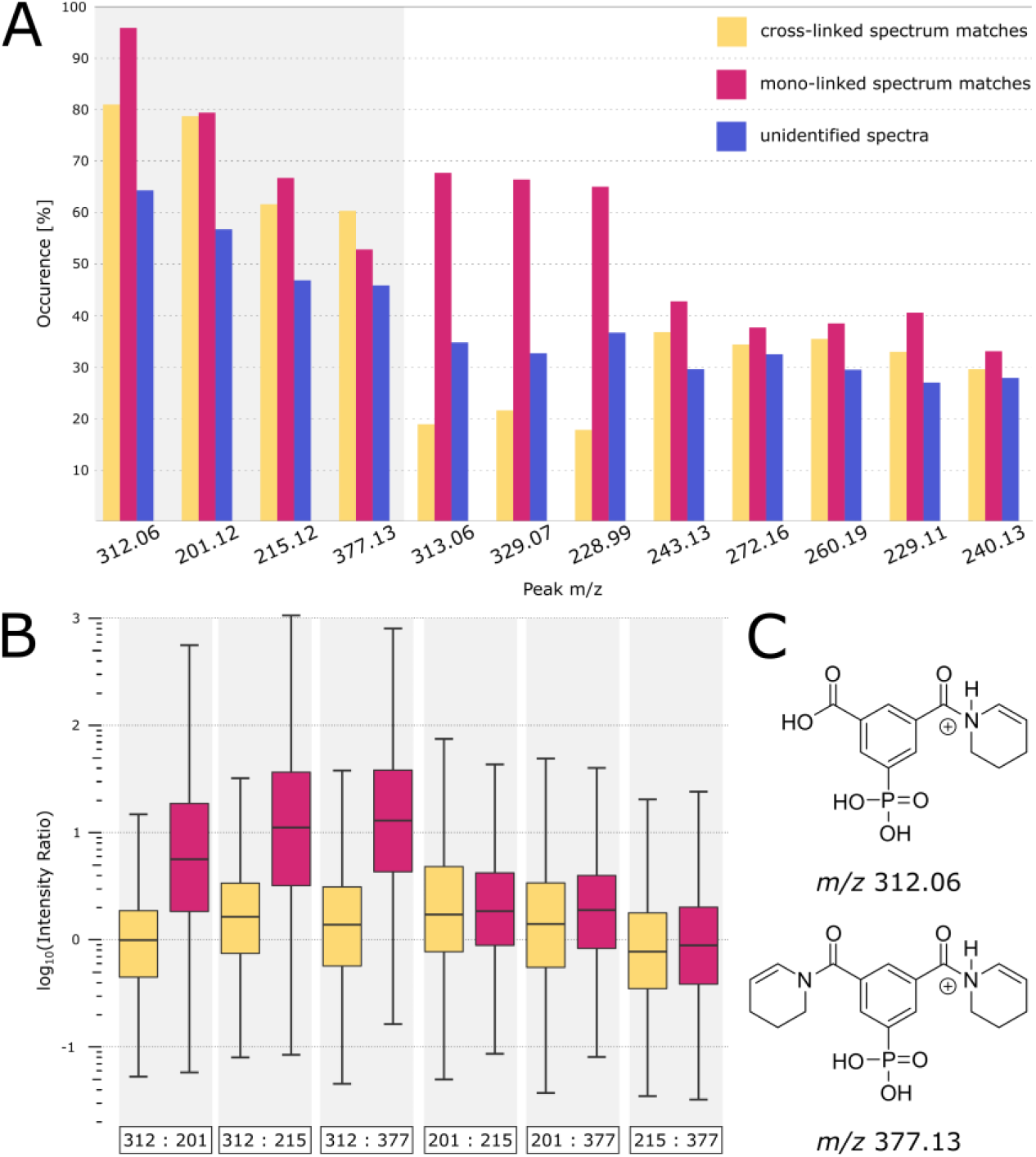
Assessment of tBu-PhoX specific diagnostic peaks and their relative intensities. (A) Representation of the most frequently occurring (in percent) peaks in cross-linked, mono-linked and unidentified spectra of a cross-linked *E. coli* sample measured with standard parameters for cross-link identification. (B) Intensity ratios of four identified diagnostic peaks to show the differences between cross-links and mono-links. (C) Proposed chemical structures of the fragmentation products that give the diagnostic peaks.

To probe the efficiency of diagnostic peak generation under different mass spectrometric conditions, we designed a method in which each MS1 precursor (with charge +3 – +8) is fragmented with a standard high-resolution MS2 scan, which is used to identify the precursor as a cross-link or a mono-link, followed by MS2 scans in which only one parameter of the standard scan is altered (Figure 2A). For parameter testing we focused on cross-linked *E. coli* as a high complexity sample to avoid bias from individual proteins. We monitored the occurrence of diagnostic peaks in the respective species and their intensity ratios. Interestingly, the intensity ratios of the diagnostic peaks are highly stable under varying MS2 scanning conditions independent of the applied fragmentation energy, orbitrap resolution, injection time and AGC target (Figure 2B). Their occurrence however increases with higher fragmentation energy (Figure 2C). This effect is most prominent for peak m/z 312.06, which can barely be detected for cross-links at low collision energy or low signal-to-noise. Considering that a survey scan that is searched against the real-time library needs to be fast to retain short duty cycles, we adapted the following scan parameters for the survey scans: scan range 100 – 500 m/z, NCE > 40%, orbitrap resolution 7.5k or ion trap rapid scan, injection time 11 – 22 ms and 50% AGC target.

**Figure 2.**
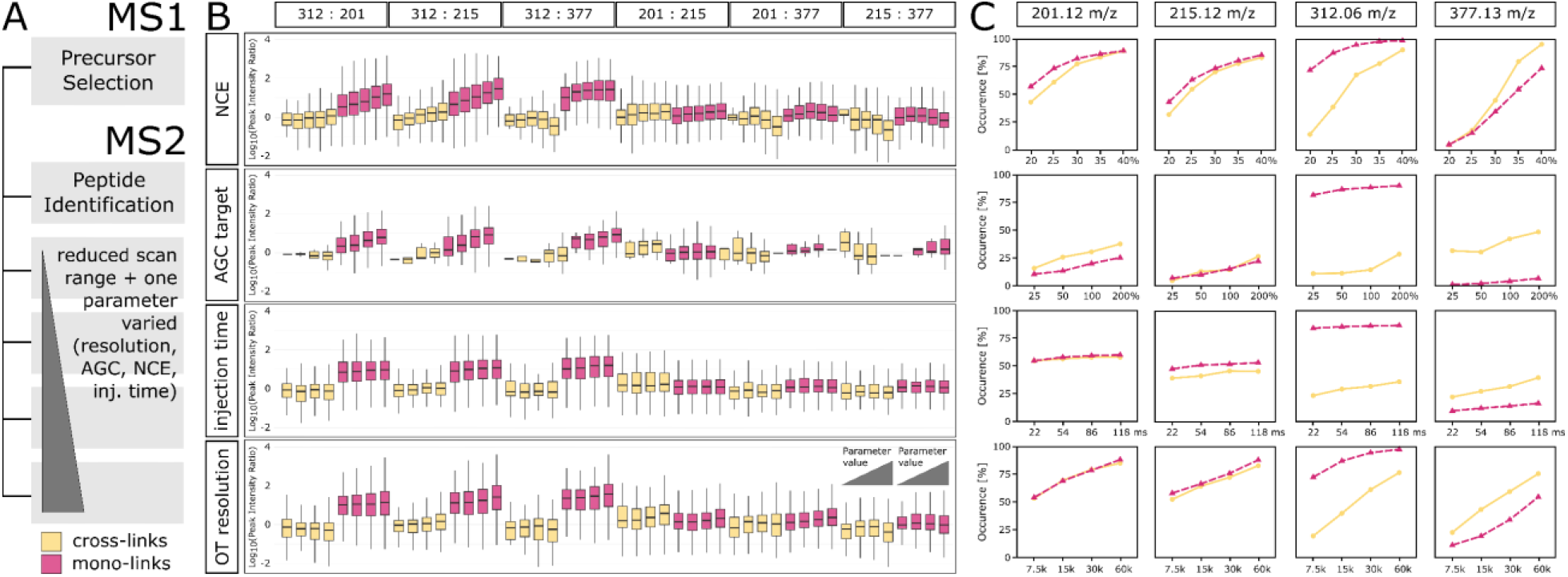
(A) Method to optimize MS parameters for the generation of diagnostic peaks. (B) Intensity ratios of the diagnostic peaks in cross-linked and mono-linked precursors using various MS2 scan parameters. NCE: 20%, 25%, 30%, 35%, 40%. AGC: 25%, 50%, 100%, 200%. Injection time: 22 ms, 54 ms, 86 ms, 118 ms. Orbitrap resolution: 7.5k, 15k, 30k, 60k. (C) Occurrence of the diagnostic peaks in cross-linked and mono-linked precursors with various MS2 parameters.

The RTLS tailored survey scans are used to fragment MS1 precursors to obtain their diagnostic peak intensity ratios which are compared to the library to mark the precursors as “potential cross-links” or “potential mono-links” (Figure 3). Only the species of interest are subjected to high-resolution MS2 scans with standard parameters.

**Figure 3.**
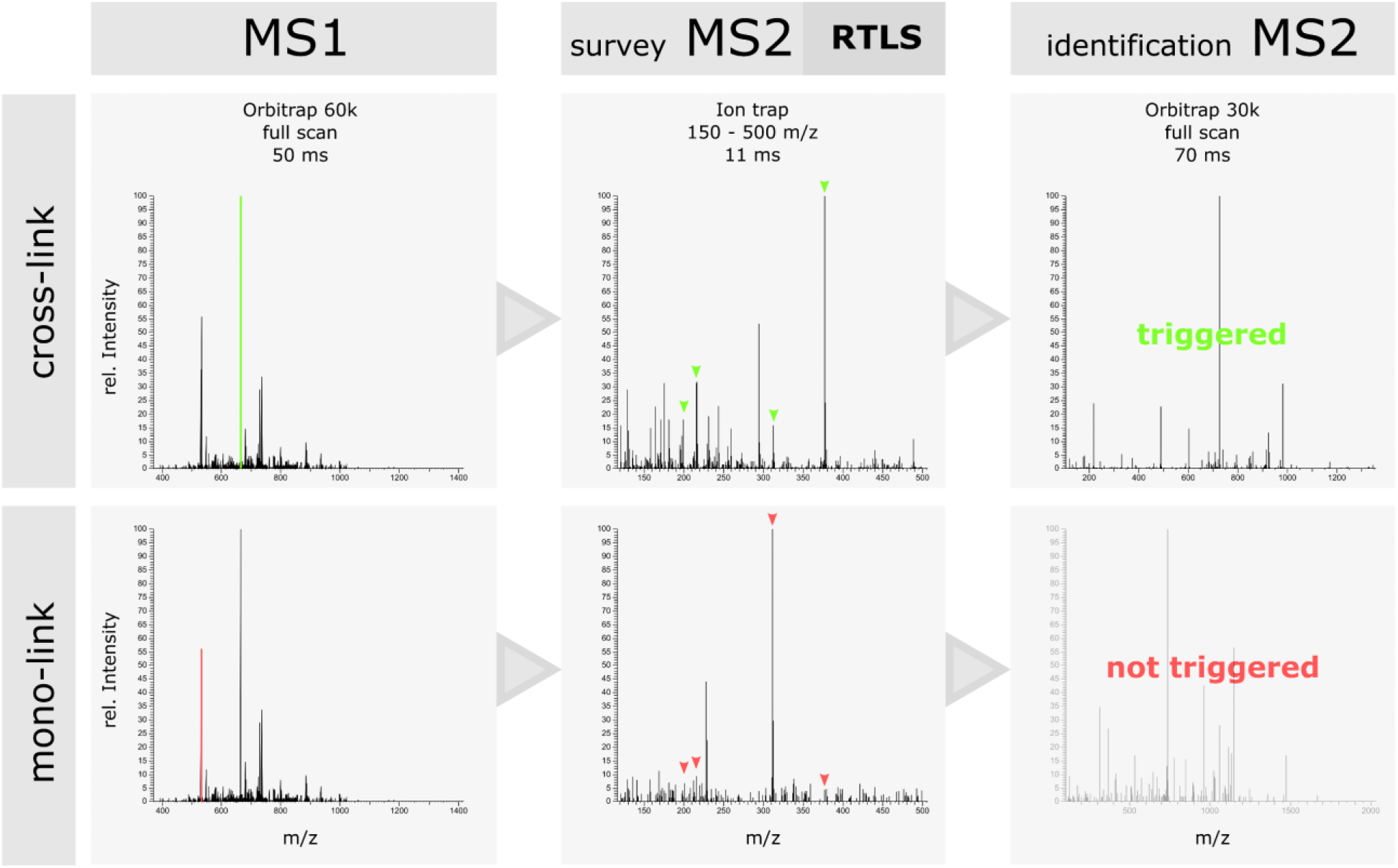
Scheme of the “cross-link promote” RTLS method. Each MS1 precursor of charge +3 – +8 is fragmented in a fast MS2 survey scan. The intensity ratios of the four diagnostic peaks are compared to a library and thereafter the precursor is labeled as a “potential cross-link” or a “potential mono-link”. A second, high-resolution MS2 scan is triggered exclusively on potential cross-links to obtain sequence information.

The quality of the match of a survey spectrum to the library is given by the cosine score of the search^15^. To increase the confidence of cross-link triggering, we tested appropriate cosine score thresholds for the survey scan in the orbitrap or ion trap from 20 to 95 (Figure 4A). Especially in the ion trap we observed that the penalty for stringency is high and therefore we suggest a relatively low score threshold of 40 to preserve a high number of potential cross-links. Optimizing the normalized collision energy (NCE) used in the survey scan showed an increase of cross-link and loop-link identifications from a tBu-PhoX cross-linked *E. coli* sample at higher energies while the mono-link identifications decreased, suggesting that higher fragmentation energies may favor the RTLS-based decision for cross-link identification (Figure 4B).

**Figure 4.**
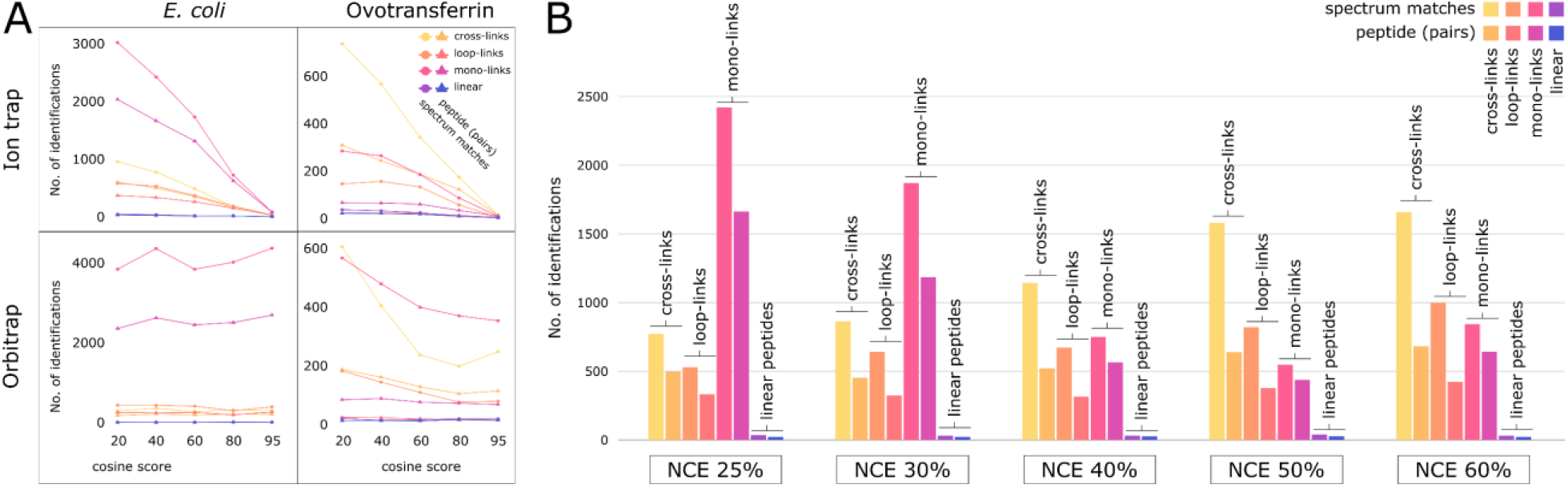
Optimizing MS parameters of the RTLS method. (A) Probing appropriate cosine score thresholds to balance confidence of the library search with preserving a high number of cross-link identifications from low- and high-complexity samples. Other MS2 survey scan parameters were kept the same. (B) Variation of the NCE in the survey scan performed with the ion trap to increase the diagnostic peak generation of cross-linked and mono-linked species.

To show the benefit of using RTLS compared to standard acquisition we monitored all identified species from a tBu-PhoX cross-linked *E. coli* sample (Figure 5A). With the “cross-link promote” and “mono-link reject” settings of the RTLS, which does not utilize a cosine score threshold, we achieved a gain in CSMs of 35% for the 60 min gradient and 14% for the 120 min gradient runs, while the mono-links were for both gradients reduced by over 80%. Interestingly, loop-linked matches also increased drastically likely due to their chemical similarity to cross-links rather than mono-links. Though the gain in CSMs in longer gradients is small, the larger improvement in the 60 min run encourages the use of shorter gradients (Figure 5A). In unenriched samples, cross-link identifications could be increased by 45% using RTLS with a short gradient (Figure 5B).

**Figure 5.**
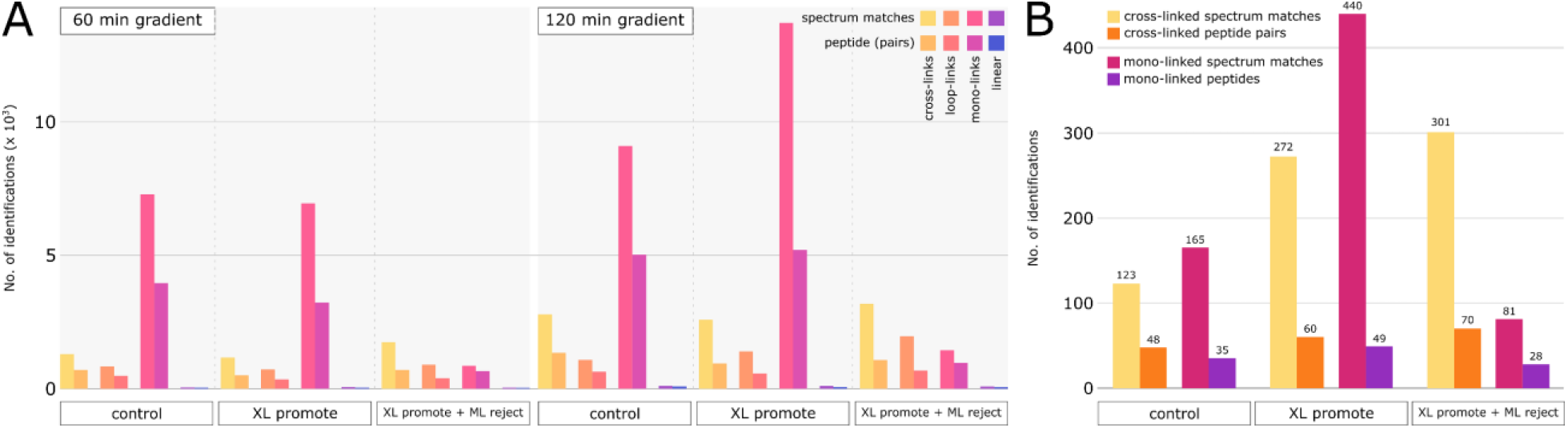
(A) Cross-link, mono-link, loop-link and linear peptide identifications from a tBu-PhoX cross-linked *E. coli* sample acquired with optimized RTLS compared to a standard MS2 acquisition strategy and two different LC gradient lengths. (B) Number of cross-links and mono-links identified from unenriched bovine serum albumin acquired with optimized RTLS compared to standard acquisition.

Next, we assessed the application of RTLS for different types of samples, i.e. from low complexity to high complexity, unenriched to enriched, and various individual proteins to avoid sequence bias. We observed RTLS is greatly beneficial for unenriched low complexity samples (Figure 5B), underscoring its advantage in samples of limited amount where cross-link enrichment is hindered. Applying RTLS to tBu-PhoX cross-linked unenriched bovine serum albumin increased cross-link identification by 45% the number of CSMs by more than two-fold. For enriched samples the advantage is lower because the number of cross-links and mono-links is almost at the maximum for low complexity samples (Figure 6, Figure S2). Furthermore, RTLS also improved the number of CSM per unique cross-link. While without RTLS most cross-links were represented by 1 – 7 CSMs, with RTLS several cross-links were found with > 10 CSMs (Figure S3). Similar results were observed when the data was searched with the XlinkX node of Proteome Discoverer 3.0 (Figure S2, Figure S4).

**Figure 6.**
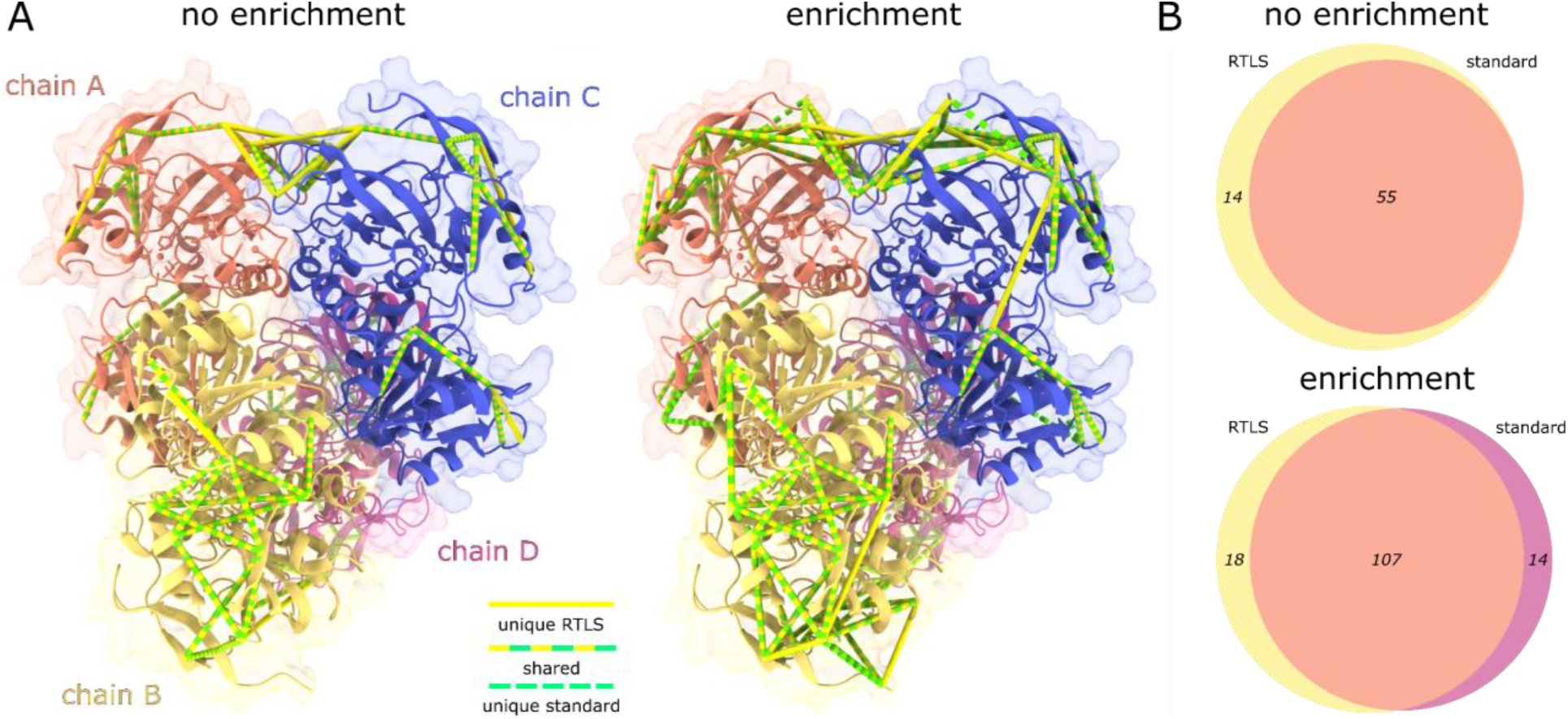
(A) Cross-link mapping of yeast alcohol dehydrogenase (PDB: 4W6Z). Cross-links identified only with RTLS (XL promote) are marked in yellow, cross-links only identified with the standard acquisition are marked in green and shared identifications are marked in yellow and green. Models are generated in ChimeraX with the XMAS plug-in^19, 20^. Only the shortest cross-links between subunits with a distance cutoff of 25 Å are displayed. (B) Venn diagrams showing the overlapped identifications between RTLS and standard measurements of the cross-links mapped in (A). Number of cross-links shown for both unenriched (upper panel) and enriched (lower panel) samples.

Furthermore, RTLS offered improvement on “cross-link resolution” for their structural analysis (Figure 6). To show the improvement of structural models obtained from RTLS measurements of enriched and unenriched samples, we acquired enriched and unenriched tBu-PhoX cross-linked yeast alcohol dehydrogenase with and without RTLS using a 60 min LC gradient. In both enriched and unenriched samples we identified additional cross-links at the oligomerization interface of the homo-tetrameric yeast alcohol dehydrogenase (PDB: 4W6Z^21^) using RTLS (Figure 6A), emphasizing the advantage of using RTLS to boost cross-link identification at specific regions and domains within the protein or protein complex of interest.

## Discussion

Recent advances in the field of XL-MS aimed to increase the number of identified cross-links from samples of different complexity^3, 9, 22^. Here, we develop a strategy that utilizes RTLS to guide cross-link acquisition by specific triggering of cross-links based on the intensity ratios of four frequently occurring diagnostic peaks (Figure 1). Two of these diagnostic peaks (m/z 201.12 and 215.12) are most likely dipeptide fragment-ions (Figure 1, Figure S1). They and other dipeptide fragment-ions (such as m/z 226.11 and 243.13) were similarly abundant in DSS samples (Figure S1), confirming their cross-linker-independent origin. In contrast, the third diagnostic peak at m/z 312.06 is specific for tBu-PhoX/PhoX mono-links. We propose that this signal is derived from a six-membered cyclic nitrogen containing ion generated from hydrolyzed mono-links (Figure 1C). This is supported by the fourth diagnostic ion at m/z 377.13, which is the most abundant signal. It is cross-link specific and also formed by the cyclization of two acylated and subsequently fragmented lysine residues^12, 13^. We demonstrate the use of these four peaks in tBu-PhoX/PhoX-modified peptide species (Figure 3), but the herein described RTLS procedure is also adaptable to other cross-linkers that generate signature fragments. Two other commonly used cross-linkers, DSS and DSSO, have been previously described to contain diagnostic peaks in their fragmentation spectra^14^.

We optimize survey scan parameters to efficiently and promptly generate diagnostic peaks for RTLS-based cross-link acquisition (Figure 4, Figure 5). Potential cross-links selected by RTLS are then triggered for MS2 identification using standard parameters. Using this acquisition strategy, we increase the number of CSMs by 35% while reducing the mono-linked spectra by more than 80% compared to standard acquisition parameters for measurements of 1 h gradients (Figure 5A, Figure S2). Specifically, the benefit of RTLS is more pronounced in unenriched samples (45% increase in cross-links) and when short gradients are in use, highlighting the advantage of RTLS to measure samples when enrichment is not possible or in high-throughput (Figure 5B). Furthermore, we show that RTLS increases the coverage of cross-links on proteins when mapping detected cross-links on high-resolution structures, and thus can provide additional information of distance constraints in cross-linking assisted structural analysis of proteins and protein complexes (Figure 6)^23^. In cases where both cross-links and mono-links are needed for structural assessment for proteins it is beneficial to perform one cross-link/loop-link dedicated acquisition with RTLS in “cross-link promote” mode and a second, mono-link focused acquisition in “mono-link promote” mode.

## Author contribution

MR, YH and RV conceived and performed the experiments. WB and RH supported the RTLS method development. DBL and JB supported the identification and characterization of diagnostic peaks. MR, YH and FL wrote the manuscript. FL supervised the research. All authors reviewed and edited the manuscript.

## Acknowledgements

The work was supported by the European Research Council (ERC) Starting Grant (ERC-STG No. 949184) and the Leibniz – Forschungsinstitut für Molekulare Pharmakologie (FMP).

## Supplementary Methods

Optimized LC-MS parameters for the RTLS-based cross-link acquisition strategy.

**Table S1.** High performance liquid chromatography peptide separation gradient.

| Retention [min] | Flow [nL/min] | Buffer B [%] |
| --- | --- | --- |
| 5 | 300 | 3 |
| 55 | 300 | 25 |
| 65 | 300 | 40 |
| 67 | 300 | 98 |
| 78 | 300 | 98 |

**Table S2.**
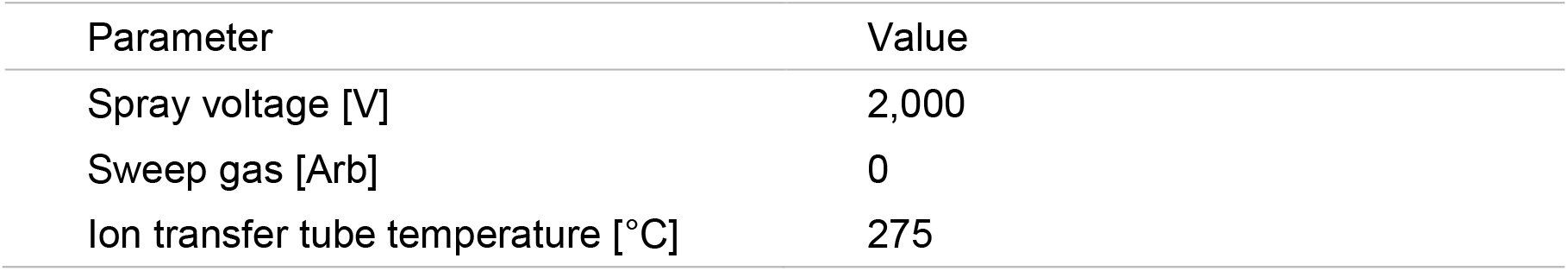
Electrospray ionization parameters.

| Parameter | Value |
| --- | --- |
| Spray voltage [V] | 2,000 |
| Sweep gas [Arb] | 0 |
| Ion transfer tube temperature [°C] | 275 |

**Table S3.**
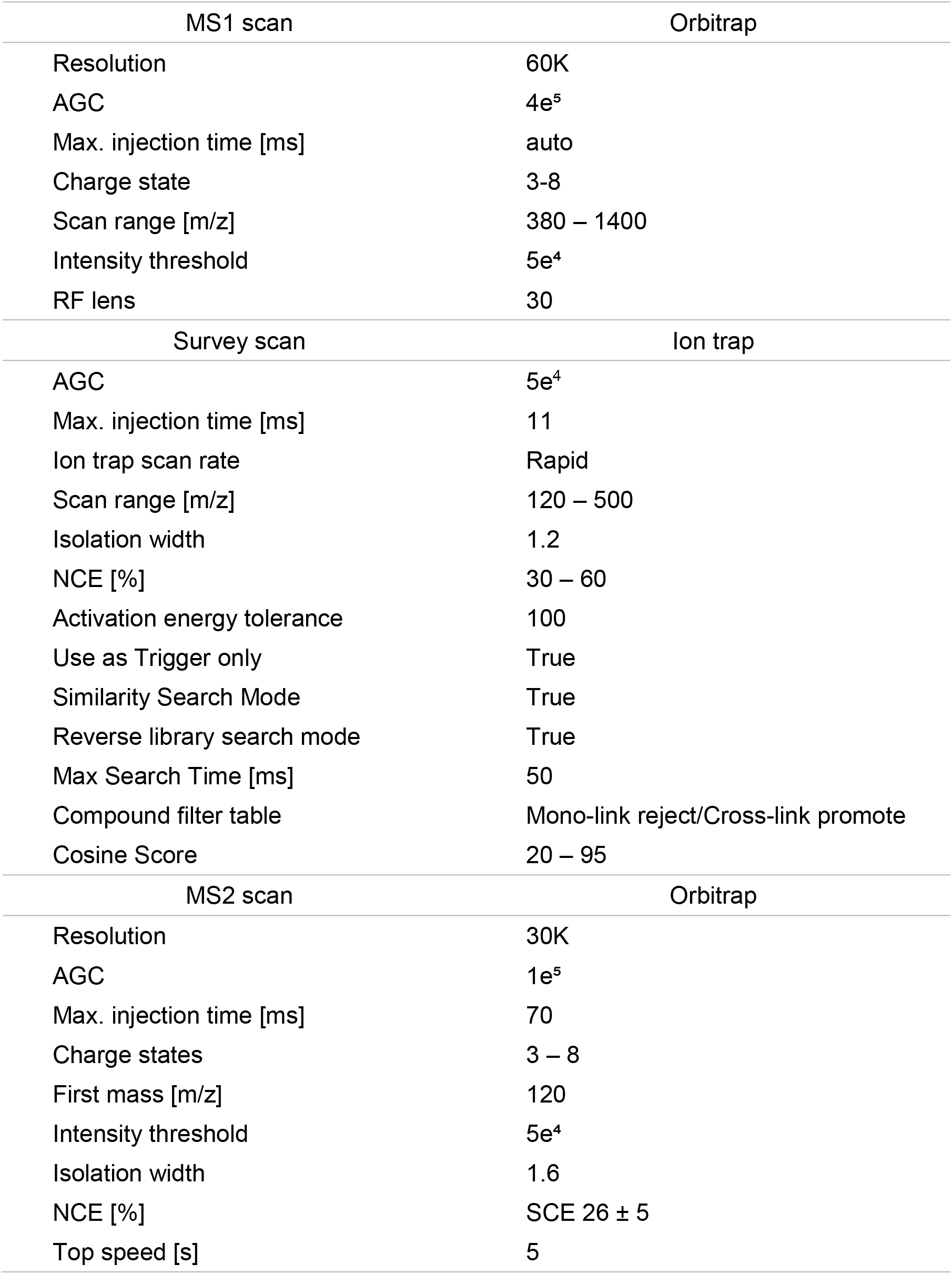
Mass spectrometric parameters for RTLS-based cross-link acquisition using an Orbitrap Eclipse.

| MS1 scan | Orbitrap |
| --- | --- |
| Resolution | 60K |
| AGC | 4e<sup>5</sup> |
| Max. injection time [ms] | auto |
| Charge state | 3-8 |
| Scan range [m/z] | 380 – 1400 |
| Intensity threshold | 5e<sup>4</sup> |
| RF lens | 30 |
| Survey scan | Ion trap |
| AGC | 5e<sup>4</sup> |
| Max. injection time [ms] | 11 |
| Ion trap scan rate | Rapid |
| Scan range [m/z] | 120 – 500 |
| Isolation width | 1.2 |
| NCE [%] | 30 – 60 |
| Activation energy tolerance | 100 |
| Use as Trigger only | True |
| Similarity Search Mode | True |
| Reverse library search mode | True |
| Max Search Time [ms] | 50 |
| Compound filter table | Mono-link reject/Cross-link promote |
| Cosine Score | 20 – 95 |
| MS2 scan | Orbitrap |
| Resolution | 30K |
| AGC | 1e<sup>5</sup> |
| Max. injection time [ms] | 70 |
| Charge states | 3 – 8 |
| First mass [m/z] | 120 |
| Intensity threshold | 5e<sup>4</sup> |
| Isolation width | 1.6 |
| NCE [%] | SCE 26 ± 5 |
| Top speed [s] | 5 |

**Figure S1.**
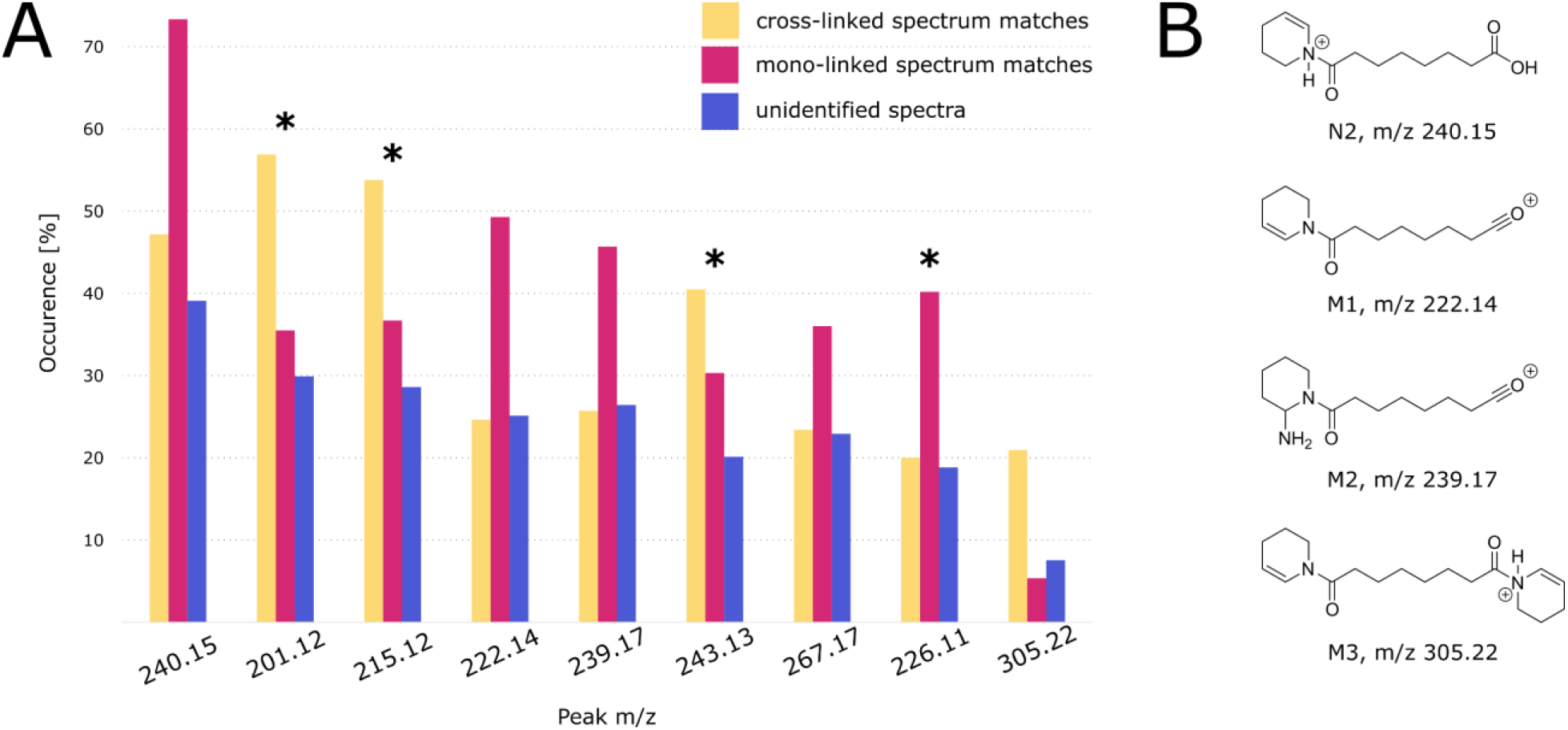
Assessment of DSS specific diagnostic peaks. (A) Representation of the most frequently occurring (in percent) peaks in cross-linked, mono-linked and unidentified spectra, measured with standard parameters for cross-link identification. Peaks that were also found with high occurrence in a tBu-PhoX cross-linked sample are marked with an asterisk. (B) Proposed chemical structures of selected fragmentation products that give diagnostic peaks.

**Figure S2.**
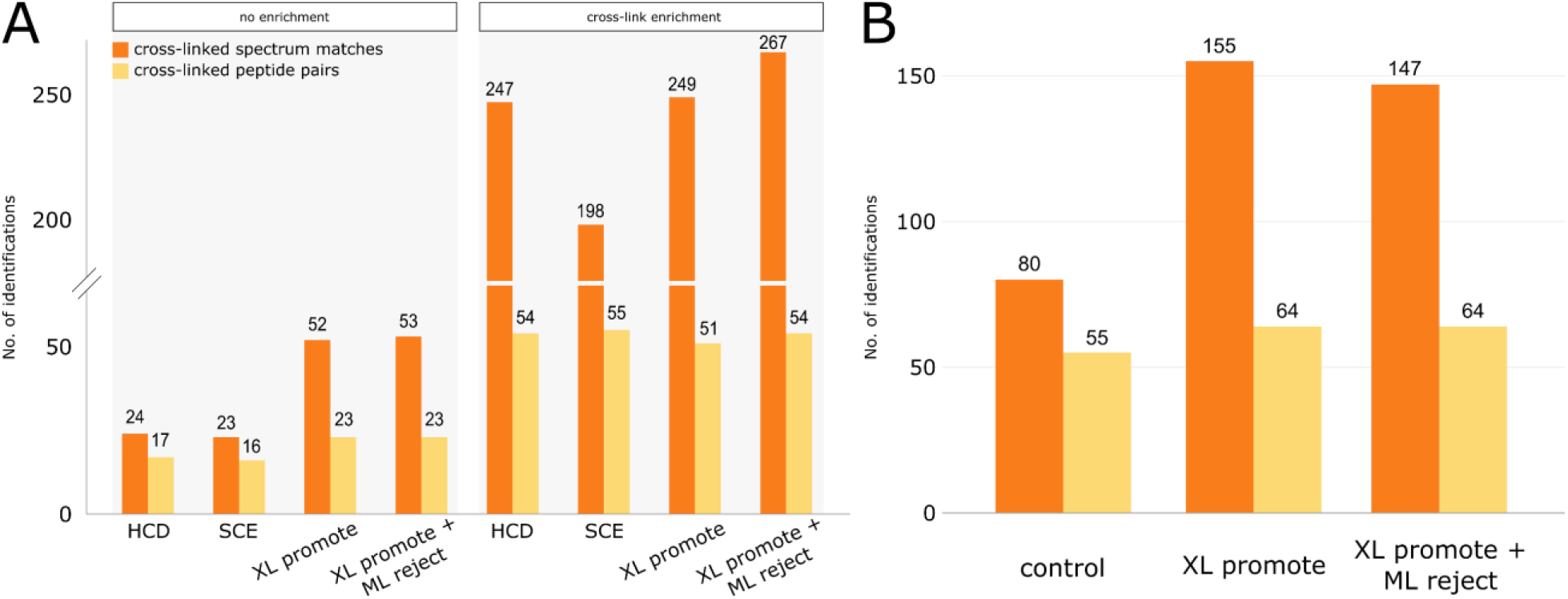
Number of CSMs and identified cross-linked peptide pairs from (A) unenriched and enriched yeast alcohol dehydrogenase and (B) unenriched bovine serum albumin using different acquisition strategies with and without RTLS. Cross-link search performed with Proteome Discoverer 3.0 XlinkX node.

**Figure S3.**
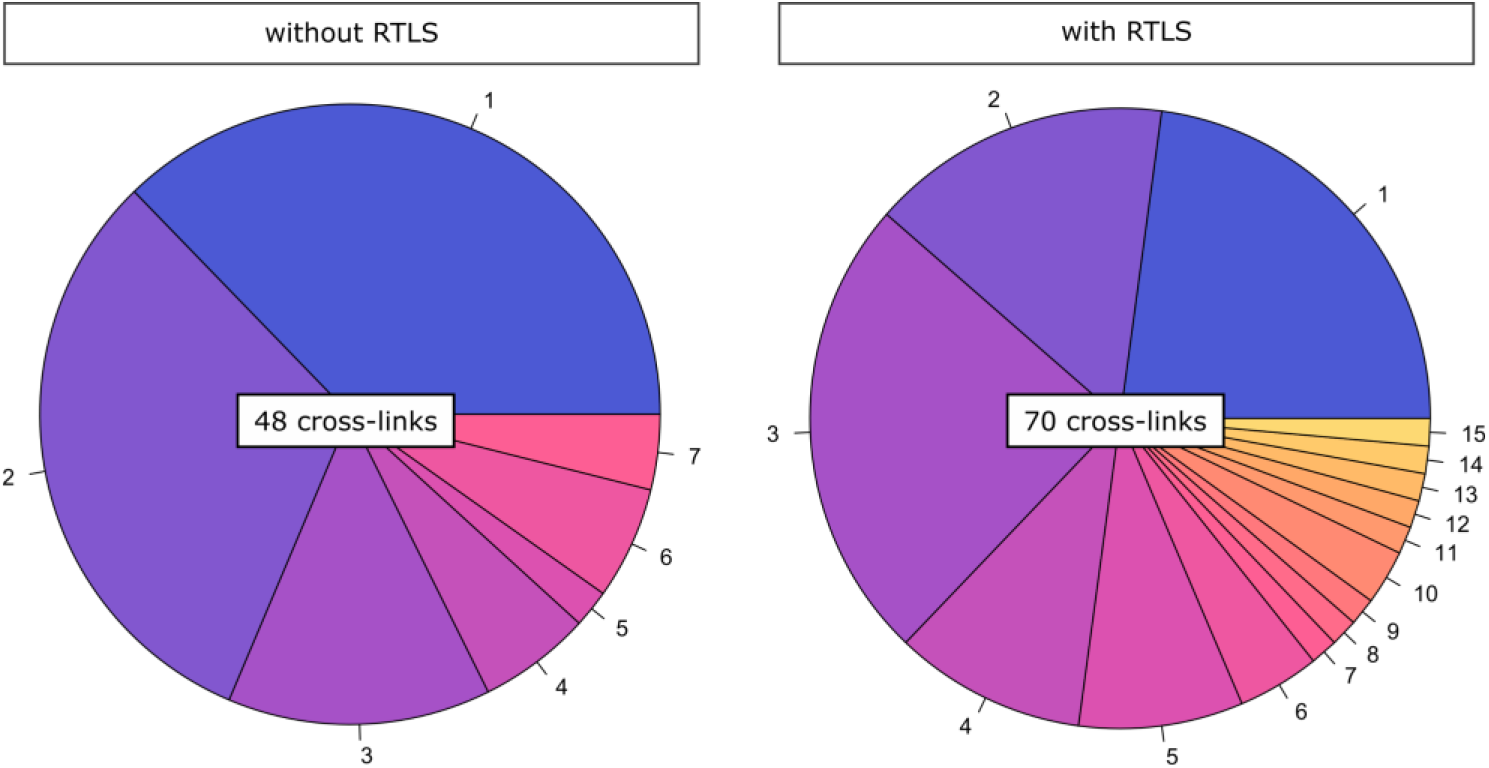
Comparison of the number of CSMs per unique cross-link identification from an unenriched bovine serum albumin sample with (70 XLs) and without (48 XLs) RTLS. Cross-link search was performed with pLink2.

**Figure S4.**
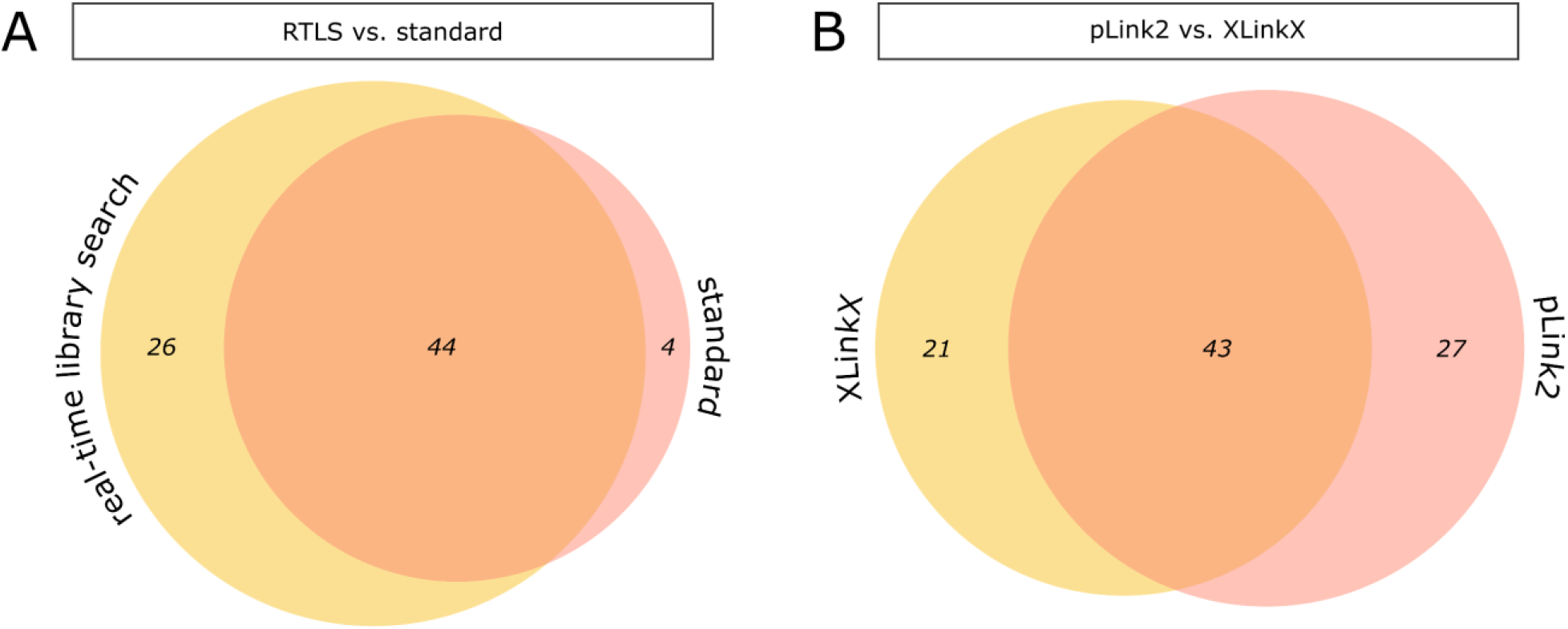
Comparing different analysis strategies in an unenriched cross-linked bovine serum albumin sample. (A) Overlap between cross-link identification from RTLS-based and standard acquisition. (B) Overlap between cross-links identified using pLink2 and Proteome Discoverer 3.0 XlinkX.

## References

1. Liu, F.; Rijkers, D. T.; Post, H.; Heck, A. J., Proteome-wide profiling of protein assemblies by cross-linking mass spectrometry. Nat Methods 2015, 12 (12), 1179–84.

2. Klykov, O.; Kopylov, M.; Carragher, B.; Heck, A. J. R.; Noble, A. J.; Scheltema, R. A., Label-free visual proteomics: Coupling MS- and EM-based approaches in structural biology. Mol Cell 2022, 82 (2), 285–303.

3. Stieger, C. E.; Doppler, P.; Mechtler, K., Optimized Fragmentation Improves the Identification of Peptides Cross-Linked by MS-Cleavable Reagents. J Proteome Res 2019, 18 (3), 1363–1370.

4. Liu, F.; Lossl, P.; Scheltema, R.; Viner, R.; Heck, A. J. R., Optimized fragmentation schemes and data analysis strategies for proteome-wide cross-link identification. Nat Commun 2017, 8, 15473.

5. Schnirch, L.; Nadler-Holly, M.; Siao, S. W.; Frese, C. K.; Viner, R.; Liu, F., Expanding the Depth and Sensitivity of Cross-Link Identification by Differential Ion Mobility Using High-Field Asymmetric Waveform Ion Mobility Spectrometry. Anal Chem 2020, 92 (15), 10495–10503.

6. Chen, Z. L.; Meng, J. M.; Cao, Y.; Yin, J. L.; Fang, R. Q.; Fan, S. B.; Liu, C.; Zeng, W. F.; Ding, Y. H.; Tan, D.; Wu, L.; Zhou, W. J.; Chi, H.; Sun, R. X.; Dong, M. Q.; He, S. M., A high-speed search engine pLink 2 with systematic evaluation for proteome-scale identification of cross-linked peptides. Nat Commun 2019, 10 (1), 3404.

7. Pirklbauer, G. J.; Stieger, C. E.; Matzinger, M.; Winkler, S.; Mechtler, K.; Dorfer, V., MS Annika: A New Cross-Linking Search Engine. J Proteome Res 2021, 20 (5), 2560–2569.

8. Matzinger, M.; Vasiu, A.; Madalinski, M.; Muller, F.; Stanek, F.; Mechtler, K., Mimicked synthetic ribosomal protein complex for benchmarking crosslinking mass spectrometry workflows. Nat Commun 2022, 13 (1), 3975.

9. Steigenberger, B.; Pieters, R. J.; Heck, A. J. R.; Scheltema, R. A., PhoX: An IMAC-Enrichable Cross-Linking Reagent. ACS Cent Sci 2019, 5 (9), 1514–1522.

10. Burke, A. M.; Kandur, W.; Novitsky, E. J.; Kaake, R. M.; Yu, C.; Kao, A.; Vellucci, D.; Huang, L.; Rychnovsky, S. D., Synthesis of two new enrichable and MS-cleavable cross-linkers to define protein-protein interactions by mass spectrometry. Org Biomol Chem 2015, 13 (17), 5030–7.

11. Piersimoni, L.; Kastritis, P. L.; Arlt, C.; Sinz, A., Cross-Linking Mass Spectrometry for Investigating Protein Conformations and Protein-Protein Interactions horizontal line A Method for All Seasons. Chem Rev 2022, 122 (8), 7500–7531.

12. Santos, L. F.; Iglesias, A. H.; Gozzo, F. C., Fragmentation features of intermolecular cross-linked peptides using N-hydroxy-succinimide esters by MALDI- and ESI-MS/MS for use in structural proteomics. J Mass Spectrom 2011, 46 (8), 742–50.

13. Iglesias, A. H.; Santos, L. F.; Gozzo, F. C., Collision-induced dissociation of Lys-Lys intramolecular crosslinked peptides. J Am Soc Mass Spectrom 2009, 20 (4), 557–66.

14. Steigenberger, B. S. H. B., Pieters R. J., Scheltema R. A., Finding and using diagnostic ions in collision induced crosslinked peptide fragmentation spectra. International Journal of Mass Spectrometry 2019, 444 (116184).

15. Bills, B.; Barshop, W. D.; Sharma, S.; Canterbury, J.; Robitaille, A. M.; Goodwin, M.; Senko, M. W.; Zabrouskov, V., Novel Real-Time Library Search Driven Data Acquisition Strategy for Identification and Characterization of Metabolites. Anal Chem 2022, 94 (9), 3749–3755.

16. Schweppe, D. K.; Eng, J. K.; Yu, Q.; Bailey, D.; Rad, R.; Navarrete-Perea, J.; Huttlin, E. L.; Erickson, B. K.; Paulo, J. A.; Gygi, S. P., Full-Featured, Real-Time Database Searching Platform Enables Fast and Accurate Multiplexed Quantitative Proteomics. J Proteome Res 2020, 19 (5), 2026–2034.

17. Jiang, P. L.; Wang, C.; Diehl, A.; Viner, R.; Etienne, C.; Nandhikonda, P.; Foster, L.; Bomgarden, R. D.; Liu, F., A Membrane-Permeable and Immobilized Metal Affinity Chromatography (IMAC) Enrichable Cross-Linking Reagent to Advance In Vivo Cross-Linking Mass Spectrometry. Angew Chem Int Ed Engl 2022, 61 (12), e202113937.

18. Kockmann, T.; Panse, C., The rawrr R Package: Direct Access to Orbitrap Data and Beyond. J Proteome Res 2021, 20 (4), 2028–2034.

19. Lagerwaard, I. M.; Albanese, P.; Jankevics, A.; Scheltema, R. A., Xlink Mapping and AnalySis (XMAS) - Smooth Integrative Modeling in ChimeraX. bioRxiv 2022.

20. Pettersen, E. F.; Goddard, T. D.; Huang, C. C.; Meng, E. C.; Couch, G. S.; Croll, T. I.; Morris, J. H.; Ferrin, T. E., UCSF ChimeraX: Structure visualization for researchers, educators, and developers. Protein Sci 2021, 30 (1), 70–82.

21. Raj, S. B.; Ramaswamy, S.; Plapp, B. V., Yeast Alcohol Dehydrogenase Structure and Catalysis. Biochemistry-Us 2014, 53 (36), 5791–5803.

22. Beveridge, R.; Stadlmann, J.; Penninger, J. M.; Mechtler, K., A synthetic peptide library for benchmarking crosslinking-mass spectrometry search engines for proteins and protein complexes. Nat Commun 2020, 11 (1), 742.

23. Klykov, O.; van der Zwaan, C.; Heck, A. J. R.; Meijer, A. B.; Scheltema, R. A., Missing regions within the molecular architecture of human fibrin clots structurally resolved by XL-MS and integrative structural modeling. Proc Natl Acad Sci U S A 2020, 117 (4), 1976–1987.

